# Beyond Pre-Post Surveys: Exploring Validity Evidence For The Use Of Experience Sampling Methods To Measure Student Anxiety In Introductory Biology

**DOI:** 10.1101/2024.06.06.597780

**Authors:** Maryrose Weatherton, Joshua M. Rosenberg, Louis Rocconi, Elisabeth E. Schussler

**Author notes:** Corresponding Author: Maryrose Weatherton, Department of Theory and Practice in Teacher Education, University of Tennessee-Knoxville, Knoxville, TN, 37996, 429 Claxton Hall, 915 Volunteer Boulevard Knoxville, TN, 37996.

## Abstract

Classroom experiences elicit many emotions in students. Of these emotions, anxiety, or the feeling of uncertainty about a prospective event, has outsized impacts on students’ academic achievement and persistence. To understand the effects of student anxiety on student outcomes, education researchers often utilize single or dual-time point (e.g., pre-post) surveys. However, we posit that these methods have significant limitations, including the potential to miss variation in student emotions across time. One methodological solution to this problem is using experience sampling methods (ESM), or short, frequent surveys that capture student experiences as they occur. This study sought to provide evidence of validity for the use of ESM to measure student anxiety in Introductory Biology courses. We compared student anxiety data gathered via ESM and traditional survey methods (i.e., pre-post surveys) in terms of 1) differences in participant recruitment, 2) students’ experience of anxiety, and 3) the predictive ability on end-of-course outcomes. Our results revealed that: 1) Compared to non-ESM participants, the ESM group has significantly fewer men and had significantly higher perceptions of instructor support, 2) Average anxiety levels measured via ESM were lower and more variable than those measured via pre-post surveys, and 3) Post-survey measures of anxiety and average ESM measures of anxiety significantly predicted students’ final grade. We discuss the implications of these results for future applications of ESM and pre-post survey methods; specifically, we propose that ESM methods may be more advantageous than pre-post surveys in cases where researchers aim to understand intra-individual processes, like emotions.

## Introduction

Student emotions are a constant in learning environments and play a vital role in the learning process (Pekrun, 1992). Both positive and negative emotions can affect student outcomes, like their attention, cognitive processes, and sense of belonging in the classroom (Respondek et al., 2017). Positive emotions, such as curiosity, excitement, and joy increase students’ engagement with curricular material and their motivation to learn (Titsworth et al., 2013). On the other hand, negative emotions like boredom, shame, and anxiety can decrease student engagement, sense of belonging, and overall academic performance (Mazer et al., 2014; Pekrun & Stephens, 2010).

One emotion that is especially relevant in undergraduate learning environments is *anxiety*, a feeling of worry or nervousness about prospective future events (Pekrun, 2006). Undergraduate students frequently report experiencing significant anxiety in their courses, a concerning trend as elevated anxiety levels are frequently associated with decreased student performance and persistence (Akgun and Ciarrochi, 2003; England et al., 2017; England et al., 2019; Center for Collegiate Mental Health, 2021). As such, a better understanding of student anxiety and its impacts may facilitate improvements in undergraduates’ academic success and retention.

Facilitating student success and retention is especially relevant within introductory courses, such as introductory Biology. These courses are not only taken by a large proportion of undergraduate students but are also prerequisites for entry into many Science, Engineering, Technology, and Mathematics (STEM) careers (AAAS, 2011). Despite this, fewer than 40% of undergraduates who initially intend to major in STEM fields complete a degree in these areas (PCAST, 2012; Hurtado et al., 2012). Introductory courses, including Biology, significantly contribute to student attrition (Rask, 2010; Fiorini et al., 2023), and students who encounter negative emotional experiences in these courses are more likely to leave STEM majors (England et al., 2017; England et al., 2019; Witt et al., 2014). In particular, students in introductory Biology courses often face high levels of anxiety, adversely affecting their academic performance and experience (Ballen et al., 2017; Schussler et al., 2021). Given this, studying anxiety is important as a potential factor in student retention in Biology.

Student anxiety and other emotional experiences in the classroom have been measured in a variety of ways. Historically, one common approach to studying student anxiety has been to use *dual-time-point surveys* (i.e., pre-post surveys), which ask participants to report their emotional state before and after an event of interest, such as a semester of a class (England et al., 2019) or even the COVID-19 global pandemic (Fruehwirth, et al., 2021). Pre-post survey methods are extremely common within disciplinary education research (Pike, 2007), although they have several limitations, especially related to studying emotional experiences, including temporal undersampling, recall bias, and time cost, among others (Zurbriggen et al., 2021; Molsa et al., 2022). For example, it may be that student anxiety increases as exam dates approach, but because of the nature of pre-post survey methods, this variation in emotion may not be adequately captured (i.e., temporal undersampling), or students may be unable to accurately recall these feelings by the next survey event (i.e., recall bias). Thus, while it is imperative for researchers to better understand students’ emotional experiences in these courses, the prevailing methods for measuring student anxiety may not fully capture the details necessary to meaningfully address students’ experiences.

One methodological answer to the limitations of pre-post surveys is the use of intensive longitudinal methods, where participants are repeatedly surveyed over a specified time period (Bolger & Laurenceau, 2013). One of the most commonly used intensive longitudinal method in education is the *Experience Sampling Method* (ESM; Zirkel et al., 2015). As an intensive longitudinal method, ESM may be able to address limitations like temporal undersampling and recall bias, thereby providing more nuanced data related to students’ emotional experiences. ESM has been successfully used to study psychological and affective constructs such as mood disorders (aan het Rot, et al., 2012), teachers’ emotions (Martinez-Sierra et al., 2019), and student engagement (Beymer et al., 2018; Xie et al., 2019). While ESM has been previously used to examine anxiety in university students (e.g., Daros et al., 2019; Goetz et al., 2020), our study sought to apply this method to measure anxiety in large introductory biology classes, understanding which is critical for retention and success of students in these classes.

Given the limited application of ESM in this context and the existing research on anxiety in this population (England et al., 2017; England et al., 2019), this study sought to establish robust validity evidence for the use of ESM that underscores its efficacy and accuracy in capturing the dynamic emotional landscape of students in large, introductory classes. Moreover, the study sought to evaluate the validity evidence of ESM relative to traditional survey techniques, thereby facilitating a comparative analysis of the insights derived from both methodologies.Overall, this study was guided by three research questions:

1. How do ESM and non-ESM participants compare in terms of a) demographic characteristics and b) pre-course perceptions of instructor support, course difficulty, and anxiety?
2. How do data gathered via ESM and pre-post surveys compare in their ability to describe students’ mean level of anxiety and its variability throughout the semester?
3. How do data gathered via ESM and pre-post surveys compare in their ability to predict end of course outcomes?

## Literature Review

### Control Value Theory of Achievement Emotions

This study utilized Pekrun’s Control-Value theory of Achievement Emotions (herein, *Control-Value theory*) as its theoretical framework. Broadly, Control-Value theory posits the antecedents and outcomes of student emotions related to an achievement-oriented task, such as completing an assignment, taking an exam, or receiving a grade in a course (Pekrun, 2006; Pekrun et al., 2023). Emotions related to these tasks are generated from individual appraisals of their level of control over a task and their subjective value for the task (see Figure 1). These emotions can be positive, such as interest, joy, pride; or negative, such as boredom, hopelessness, and anxiety. Students’ perceptions of control and value are jointly referred to as ‘appraisals,’ and Pekrun posits that these appraisals are influenced by students’ goals and beliefs (i.e., internal factors) as well as by the design of learning and social environments (i.e., external factors), such as the classroom context, pedagogy, and consequences of failure (Figure 1). Consequently, these emotional experiences impact students’ thinking and performance.

Control-Value theory posits that emotional experiences can affect a range of outcomes, including cognition, motivation, achievement, well-being, and social relationships (Pekrun et al., 2023). Emotional experiences affect working memory, and previous work has revealed that positive emotions generally improve working memory, while negative emotions reduce working memory and information processing (Pekrun et al., 2002; Grossberg, 2009; Wijbenga et al., 2024). Previous research has revealed that students with higher levels of anxiety across a semester of introductory Biology tend to have worse grades (Ballen et al., 2017), and potential mechanisms of this relationship may include the effect of negative emotions on working memory and information processing. Given the impact of emotional experiences on students’ outcomes, it is important to further understand how these emotions vary over time in order to create more effectively-timed intervention strategies to address negative emotions in the classroom.

### Student Anxiety in Introductory Biology Courses

While Control-Value theory can be used to understand a wide range of achievement emotions, this study focused specifically on student anxiety in introductory biology courses. Anxiety is a prospective emotion that is generated when an individual appraises a task as being important, but is uncertain about the level of control they have to achieve the task (Miceli & Castelfranchi, 2005). Like all other achievement emotions, anxiety is driven by cognitive appraisals and environmental context, and has impacts on numerous outcomes. While moderate levels of anxiety can positively affect student outcomes (Yerkes & Dodson, 1908), research in the field of STEM education has come to a broad consensus about the negative impacts of high anxiety on the student experience in introductory biology (England et al., 2017; England et al., 2019; 2021a; Ballen et al., 2017; Cotner et al., 2020; Downing et al., 2020).

Previous work related to student anxiety has found that student perceptions of instructor support are highly correlated with student anxiety (Schussler et al., 2021), suggesting that students who feel more supported by their instructors may have higher perceptions of control, and thus lower anxiety. However, student-level anxiety has been found to differ within course sections taught by the same instructor, indicating that other contextual factors beyond the instructor or course structure may play a key role in student emotion (England et al., 2017). One notable contextual factor is the demographic make-up of a class, as certain groups of students often have higher levels of anxiety than others (Pekrun & Perry, 2014). For example, England and colleagues (2019) revealed that women and students who took fewer advanced biology classes in high school had significantly higher anxiety than their classmates; and the link between these demographic characteristics and anxiety is supported by numerous other studies (e.g., Ballen et al., 2017; Misra & McKean, 2000). In terms of the impacts of anxiety in introductory Biology, high levels of anxiety have been empirically linked to myriad negative outcomes, like a lower intention to persist in a Biology major (England et al., 2017; England et al., 2019; Brownlow et al., 2000), lower academic achievement in the course (Ballen et al., 2017), and lower academic self-efficacy (Barthelemy et al., 2015).

### Measurement of Student Emotion & Limitations of Pre-post Survey Methods

In his 2006 review of Control-Value theory and extant methodological gaps in the study of emotion, Pekrun argued that a significant oversight in previous research has been the treatment of emotion as a static phenomenon (Pekrun, 2006). This has led to data collection methods that overlook the dynamic and often non-linear trajectories of emotional appraisals suggested by Control-Value theory. Instead, Pekrun highlights the necessity of recognizing emotion as a complex, event-dependent phenomenon and advocates for more nuanced analyses of both intra– and inter-individual differences in emotional experiences (Pekrun, 2006). Despite this call for methodological innovation, the measurement of student emotion in introductory biology courses still predominantly relies on traditional approaches, like pre-post surveys (England et al., 2019; Pike, 2007). These methods are often chosen for their simplicity and practicality; however, they exhibit significant shortcomings in accurately measuring student emotion. Key limitations include temporal undersampling, recall bias, and data analysis challenges related to intra-individual processing. Below, how each of these may be limiting our understanding of student emotion is discussed.

Pre-post survey methods are notably impacted by temporal undersampling, which occurs when an insufficient number of surveys are conducted to accurately track changes over time in the targeted outcome. Student emotions, including anxiety, are highly dynamic across time, and as such, the use of single– or dual-time point surveys for analyzing student emotions may not capture enough variation to adequately understand the nuance in student emotions and their change across time (Pekrun, 2006).

Even if pre-post methods did capture adequate levels of variation, however, responses may still be subject to recall bias. Recall bias happens when participants are unable to accurately or completely recall their experiences; the likelihood of recall bias increases as time goes on and is impacted by participants’ current emotional states (Sedgewick, 2012; Colombo et al., 2020; Glazier & Alden, 2017). Data collected via traditional survey methods, which can ask participants to recall information after an entire semester or longer, may be unable to capture variation in anxiety that happens throughout the semester and may additionally suffer from participants’ ability to accurately recall their emotional states.

Finally, pre-post survey methods often yield too few data points to analyze individual responses in depth, necessitating a reliance on inter-individual comparisons. For example, while students with higher overall anxiety might generally perform worse on exams than their peers (an *inter*-individual finding), a single student’s anxiety levels may also vary from one exam to another (*intra*-individual variation) and this can also affect their performance (Pekrun, 2006; Schmitz & Skinner, 1993). This limitation is particularly relevant within affective research, because emotional responses and their outcomes are greatly influenced by individual factors (Hamann & Canli, 2004; Kuppens et al., 2009). As such, the reliance on interindividual covariation to investigate intraindividual psychological dynamics is a significant concern, as these methods can lead to misleading conclusions (Pekrun, 2006; Tempelaar & Niculescu, 2022; Zhang, 2022;). Cross-sectional, longitudinal data collection methods can better explore the nuances of intra-individual emotional variation, as well as the distribution of these individual differences within a larger sample (Pekrun, 2006; Pekrun & Hofmann, 1996).

Thus, prevailing methods centered on pre-post survey designs result in data that likely simplify the complex emotional environments that students are navigating. These methods are likely unable to capture variation in anxiety that may be meaningfully related to course outcomes. One way to address these gaps is utilizing longitudinal methods to study emotion; longitudinal methods capture real-time data, as opposed to retrospective reflections on emotions, and can be used to examine both intra– and inter-individual variation across time (Goetz et al., 2010; Lu et al., 2023; Xie et al., 2019, 2023). One type of longitudinal method, experience sampling methods, may be well-suited to capture these nuances within an introductory Biology course.

### The Experience Sampling Method (ESM)

ESM is an intensive longitudinal method that allows researchers to collect data about the daily life of individuals as it is perceived in the moment, at an increased level of granularity compared with traditional forms of survey research (Hektner et al, 2007; Zirkel et al., 2015). Intensive longitudinal methods is an umbrella term used to describe survey research methods that involve a) repeatedly prompting study participants to b) respond to brief questions (Bolger & Laurenceau, 2013). This description leaves room for several flavors (e.g., Experience Sampling, Ecological Momentary Assessment, Diary studies) of intensive longitudinal methods that have been developed in varied research fields (Bolger & Laurenceau, 2013). ESM combines the ecological validity of naturalistic observation, the non-intrusive nature of diaries, and the precision of scaled survey items. Researchers can observe participants’ experiences through ESM repeated measures at a level of detail equivalent to qualitative research, but at a scale typically infeasible in most qualitative research studies. We briefly expand on the affordances and drawbacks of ESM, in general, to set the stage for their use in the present study.

### Affordances of ESM

Intensive longitudinal methods are employed for a multitude of reasons. First, intensive longitudinal methods enable researchers to understand individuals’ experiences in their natural environments, capturing experiences as they occur. What people say they are feeling, thinking, or doing in the moment may differ from their retrospective accounts, which is particularly relevant in the study of emotions.

Moreover, many constructs as they are conceptualized vary dynamically across contexts and over time. For instance, theoretical accounts of student interest describe it as “the psychological state of engaging or the predisposition to reengage with particular classes of objects, events, or ideas over time” (Hidi & Renninger, 2006, p. 112). This temporal and contextual variation is not exclusive to interest but extends to many psychological constructs such as engagement, confusion, enjoyment, and anxiety (Beymer et al., 2018, 2021; Goetz et al., 2016; Inkinen et al., 2020; Sinatra et al., 2015; Schernoff & Schmidt, 2008; Schmidt et al., 2018). Traditional survey methods often fail to capture this variability, but intensive longitudinal methods can, when used deliberately, can accurately capture this variability. As Bolger and Laurenceau (2013) describe it “traditional longitudinal designs can also examine temporal unfolding, they are often limited by few repeated measurements taken over long time intervals.” (p. 5).

Another reason for the use of intensive longitudinal methods is to explore within-person relations between constructs, such as between students’ emotions (affect) and their engagement (Beymer et al., 2018). Such relations may be difficult, if not impossible, to examine using traditional survey research methods given the limited number of time points at which such relations could be examined–or, such relations may improperly be assessed using between-person data (Bolger & Laurenceau, 2013).

### Drawbacks of ESM

While intensive longitudinal methods have several benefits, their use also involves trade-offs. They require more time and effort on the part of participants, which may create a need for greater incentives. This can be important for addressing selection biases relating to who participates in such studies and who responds to ESM prompts, an issue addressed in past research (Scollon et al, 2003). These studies can also be expensive to manage; ESM data are often collected using technological tools or services that are not typically a part of university data collection software, requiring additional software expenses or expertise. Last, the data collected is often highly complex, requiring ample time for both data cleaning and analysis.

### Standards for Validity

This study sought to provide validity evidence for the use of ESM in large introductory classes, and to compare validity evidence between ESM and standard pre-post survey methods, with an overall goal to consider when and for what purposes one may use ESM versus pre-post survey methods. Given these goals, this study used the approach to validity outlined by the *Standards for Educational and Psychological Testing*, a document collaboratively developed by the American Educational Research Association (AERA), the American Psychological Association (APA), and the National Council on Measurement in Education (NCME). The *Standards* sets forth rigorous guidelines for the development and evaluation of testing and assessment practices (AERA, APA, & NCME, 2014). Central to these guidelines is the emphasis on presenting validity evidence for any assessment method used. Validity evidence is foundational, as it confirms that the test measures what it intends to measure, thereby ensuring the accuracy, fairness, and overall quality of an assessment. The 2014 *Standards* outlines 26 specific standards for validity, which are clustered into three main categories: establishing intended uses and interpretations, issues regarding samples and settings used in validation, and specific forms of validity evidence (AERA, APA, & NCME, 2014). Notably, these standards are not meant to be a checklist, but should instead be used as a frame of reference, with the applicability of each standard to be determined by the research team and the context within which the tool is to be used (AERA, APA, & NCME, 2014, pp. 7).

This paper focuses on providing validity evidence for ESM within a novel context: assessing student emotions in large, introductory biology classes. As outlined by the 2014 *Standards*, we began by establishing the intended uses and interpretations of the method, and then moved to provide specific forms of validity evidence, aligned with our research questions (Table 1). Table 1 provides a rationale for each research question and illustrates its alignment with a standard from the 2014 *Standards* document. Through this research, we seek to underscore the importance of methodological rigor and the potential of ESM to offer insights into student emotions, thus enriching the educational research landscape and enhancing our understanding of student experiences in large, introductory classes. By providing validity evidence for the use of ESM in relation to pre-post surveys in the same population, we can not only provide evidence of its efficacy and accuracy, but also suggest when each method may be most useful for measuring anxiety given the desired research outcomes.

**Table 1:** Research questions aligned with specific forms of validity evidence outlined in the 2014 Standards.

| *Research question* | *Standard* | *Aligned Rationale* |
| --- | --- | --- |
| 1: How do ESM and non-ESM participants compare in terms of demographic characteristics and pre-course perceptions? | **Standard 1.8:**<br>Describe the composition of any sample in as much detail as possible | As ESM participants are intended to represent the larger population of introductory Biology students at this university, it is imperative that this sample be described and any limitations in representativeness noted. Thus, we will evaluate the extent to which our ESM population is similar to non-ESM participants. |
| 2: How do data gathered via ESM and pre-post surveys compare in their ability to describe students' mean level of anxiety and its variability throughout the semester? | **Standard 1.14:**<br>Provide rationale and evidence to support the interpretation of scores, subscores, or score differences | The research question directly addresses Standard 1.14 by evaluating the effectiveness of different methods in describing students' anxiety experiences. This comparison allows for evidence-based interpretation of scores, providing a rationale for selecting and justifying the preferred method based on its ability to capture the nuanced aspects of anxiety among students accurately. |
| 3: How do data gathered via ESM and | **Standard 1.17:**<br>If validation relies on | By evaluating how well the data from each method correlates with final |
| pre-post surveys compare in their ability to predict end of course outcomes? | evidence that scores are related to one or more criterion variables, evidence of the suitability and quality of criteria. | grades, we aim to provide evidence of the suitability and quality of criteria used for validation purposes. By assessing their ability to predict academic performance, the study directly addresses the standard's requirement for evidence-based validation of measurement tools in relation to relevant and well-defined criterion variables. |

## Methods

### Sample

Before collecting data or contacting participants, researchers obtained approval for this study from our institution’s Institutional Review Board (*IRB Number removed for blinding*). Data for this study were collected from participants in three large introductory Biology courses (three sections of organismal biology, two sections of cellular biology, and four sections of non-majors introductory Biology) at one research-intensive university in the Southeastern United States. Data were collected during the spring semester of 2022. There were a total of 1,884 students enrolled in these courses.

Data for this study came from three sources: 1) a pre-survey, given at week one of the semester, 2) ESM data collected throughout the semester (weeks three through fifteen), 3) a post-survey given at week fourteen of the semester. Our sample size varied across these measures. At week one, 1,014 students completed the pre-survey, and 941 students completed the post-survey at week fourteen. 278 students (27.1% of the students who completed the pre-survey) agreed to participate in the ESM portion of the study. The 278 students who participated in the intensive longitudinal method portion of the study completed a total of 6,016 surveys (*M* = 21.6 per student; *SD* = 9.02; *max.* = 30; *min.* = 1). 176 students completed the pre-survey, post-survey, and responded to three or more ESM prompts. We decided to exclude students who responded to three or fewer ESM prompts, as we reasoned that this was a principled cut-off given that ESM is predicated on the additional data points the approach can yield, relative to pre-post surveys. Students meeting these standards were herein considered ‘complete observations’ and were the sample used as data for this study (*N* = 176).

## Data Sources and Procedure

### Pre-Survey

At week one of the semester, instructors invited students to participate in a survey by distributing recruitment emails from the researchers. Students were incentivized to take part in the pre-course survey with a small number of bonus points toward their overall course grade (up to 4 points out of a total of approximately 1,000 course points, with the number of points determined by the individual instructor).

The pre-course survey was distributed via Qualtrics and collected data about students’ self-reported demographic information, their level of anxiety (four items), their perception of course difficulty (three items), their perception of instructor support and whether they would be interested in participating in ESM data collection throughout the semester (Supplementary Material, Appendix A). Specific questions about student anxiety, perceptions of instructor support, perception of difficulty, and self-reported demographic information were adapted from previous surveys used in the same context (for more information, see England et al., 2019; Schussler et al., 2021). England et al. (2019) provided evidence of the validity of these questions within the course and student populations, including adequate factor loadings within exploratory and confirmatory factor analysis (England et al., 2019; Weatherton et al.; *in review*). Students who agreed to participate in ESM data collection provided their phone number, student identification number, and the days and times of their biology lectures. These data were entered into our ESM data collection application described next.

### ESM Data Collection: Short Message Survey

We used an ESM data collection app, [name blinded for review] (Rosenberg & Lishinski, 2021), that we developed instead of using a third-party platform. We believed this approach would minimize the risk of self-selection bias and respondent fatigue compared to other app-based ESM data collection systems. App-based systems rely on students interacting with an unfamiliar interface, which has the disadvantages of lower ease of use as well as an increased perception of obtrusiveness compared to the SMS (i.e., text message) interface that students are already familiar with. By leveraging such an interface for our data collection, we were able to minimize the burden on students when responding to our items, as well as more seamlessly integrate our surveys into their everyday lives. Beyond the benefits of optimizing the user experience for students responding to our ESM surveys, the app also offered the benefit of full customizability to meet the needs of our data collection procedures in order to reach each individual student at the optimal time.

Data collection was conducted by using the app to send a prompt to students a few minutes after their Biology lecture session had ended. We sent participants one survey question after each class. This survey question was one of four items from the measure of anxiety used in the pre-survey (Figure 2). These items were randomly rotated, so students were prompted with a different item after each class. ESM data collection began February 7th, 2022 (week three of the spring semester) and continued until May 9th, 2022 (week fifteen of the spring semester). On average, each survey typically took students less than 30 seconds to complete. Participants were sent prompts up to 30 times over the course of the semester, and the compensation structure was such that participants earned additional compensation for every survey they completed, for a potential total compensation of $30 for participating.

On average, students from the ‘complete responses’ group (*<u>N</u>* = 176) responded to 26.35 (*SD* = 4.59) items. Figure 3 shows nine individual student responses as an example of the variability in response numbers, patterns across students, and anxiety trends over time. For example, student 9 reported relatively high anxiety at the beginning of the semester, which increased around mid-March and then decreased through the rest of the semester.

### Post-Survey

At week 14 of the course (out of a total of 16 weeks), instructors again invited all students to participate in a survey by distributing recruitment emails from researchers. As with the pre-survey, participation was incentivized via a small number of bonus points towards students’ overall course grade. The survey (herein referred to as ‘post-survey’) was structured identically to the pre-survey, with the exception of the section where participants could sign up to participate in ESM data collection.

### Course Grades

After the course was completed, instructors provided researchers with the final course grades for all students who consented to participate in the research project and consented to have their grades shared. The instructors shared the percentage of points earned out of total points possible to earn (as a value ranging from 0-100) for students’ final grade. Table 2 provides an overview of the variables used in this study. In subsequent sections, we refer only to these variable names for consistency and clarity.

**Table 2:** Variables in the study.

| **Variable Name** | **Variable Type** | **Item(s)** | **Data Source(s)** | **Corresponding RQ(s)** |
| --- | --- | --- | --- | --- |
| Gender identity | Categorical | What is your gender identity? | Pre-course survey | 1a |
| Racial identity | Categorical | What is your racial/ethnic identity? | Pre-course survey | 1a |
| Year in school | Categorical | What year are you in school? | Pre-course survey | 1a |
| Time since last biology class | Categorical | How long has it been since you last took a biology class? | Pre-course survey | 1a |
| Course | Categorical | What is the name of the professor of your Biology lecture class this semester? | Pre-course survey | 1a |
| Perception of instructor support | Ordinal (1-10) | On a 10-point scale, how supportive do you feel the instructor of your biology class has been so far? | Pre-course survey | 1b |
| Perception of course difficulty | Continuous, (Mean of all items, Likert-type scale 1-7) | 1: Biology lecture is complex<br>2: Biology lecture is complicated<br>3: Biology lecture is difficult | Pre-course survey | 1b |
| Pre-survey anxiety | Continuous (Mean of all items, Likert-type scale 1-7) | 1: Biology lecture makes me nervous<br>2: Biology lecture is stressful<br>3: Biology lecture makes me anxious<br>4: Biology lecture scares me | Pre-course survey | 1b, 3 |
| Post-survey anxiety | Continuous (Mean of all items, Likert-type | 1: Biology lecture makes me nervous | Post-course survey | 3 |
|  | scale 1-7) | 2: Biology lecture is stressful<br>3: Biology lecture makes me anxious<br>4: Biology lecture scares me |  |  |
| Mean ESM anxiety score | Continuous, mean of all ESM responses per student (Likert-type scale, 1-7) | 1: Biology lecture makes me nervous<br>2: Biology lecture is stressful<br>3: Biology lecture makes me anxious<br>4: Biology lecture scares me | ESM data collection | 2, 3 |
| Mean pre-post anxiety score | Continuous, calculated by adding the pre-score to the post-score and then dividing the sum by two. | N/A | Pre-course survey<br>Post-course survey | 2 |
| Pre-post | Continuous, | N/A | Pre-course | 2 |
| change score | calculated by subtracting pre-survey anxiety score from post-survey anxiety score. |  | survey<br>Post-course<br>survey |  |
| Final grade | Continuous, 0-100 | N/A | Sent by<br>course<br>instructor<br>after course<br>had ended | 3 |

## Data Analyses

All of our analyses were conducted in R (R Core Team, 2024) using the following packages: MASS (Venables & Ripley, 2002), tidyverse (Wickham et al., 2019), sjPlot (Lüdecke, 2023), and lme4 (Bates et al., 2015).

### RQ 1a: How do ESM participants compare to non-ESM participants in terms of demographic characteristics?

We compared demographic data between students who elected to participate in ESM data collection (N = 176) and those who did not (i.e., those who completed the pre-course survey but did not enroll in ESM data collection, N = 832). We evaluated differences in demographic factors using the following data collected during the pre-survey: self-reported racial and ethnic identity, self-reported gender identity, year in school, course, and time since last biology class. We used Chi-square tests to evaluate whether any differences between the two groups were statistically significant (Franke et al., 2012).

### RQ 1b: How do ESM participants compare to non-ESM participants in terms of pre-course perceptions?

We compared differences in the student perceptions using the following data collected during the pre-survey: students’ level of anxiety related to the course, course difficulty, and instructor support. We used Wilcoxon signed-rank tests to compare the mean perception scores between ESM and non-ESM participants to determine if there were statistically significant differences in their perceptions of anxiety, course difficulty, or instructor support. This non-parametric approach was selected due to the non-normal distribution of the data.

### RQ 2: How do data gathered via ESM and pre-post surveys compare in their ability to describe students’ mean level of anxiety and its variability throughout the semester?

To examine general trends in student anxiety over time for the 176 participants, we calculated and compared the mean and standard deviation of anxiety measured via ESM and pre-post surveys. We performed a Shapiro-Wilk test for normality and found that the difference between students’ average ESM and pre-post survey scores were normally distributed (*W* = 0.989, *p* = 0.167). Thus, we used paired *t*-tests to test for differences between the two groups (Hsu & Lachenbruch, 2014).

Furthermore, because participants responded to multiple ESM prompts during the study, we were able to examine the variation in their anxiety over time. Two students may have had a similar mean ESM anxiety, but their level of variation around that mean may have differed (e.g. an average ESM anxiety of four with a range in variation from one-seven has a standard deviation of three, versus a range in variation from three-five having a standard deviation of 0.25), adding further nuance to our understanding of their experience. Thus, we also calculated the mean level of intra-student variation in ESM responses. Given that the pre-post survey measures yielded only two data points per participant, the usefulness and interpretability of the standard deviation in such a case is limited. Instead, to gauge variation, we calculated change scores by subtracting post-survey anxiety from pre-survey anxiety score for each student.

### RQ 3: How do data gathered via ESM and pre-post surveys compare in their ability to predict end of course outcomes?

To evaluate the predictive ability of the two measures of anxiety (i.e., ESM and pre-post surveys) on students’ final course grade, we compared the output of three separate linear models with the same outcome variable, final grade. Each model had a nearly-identical set of predictors, save for the measure of anxiety (see regression equation). Model A used students’ pre-survey anxiety scores as the measure of anxiety, Model B used students’ post-survey anxiety scores, and Model C used students’ mean ESM anxiety scores as the measure of anxiety.

Grade = β0 +β1 * **Anxiety** + β2 *instructor + β3*course + β4*time since last class + β5*racial identity + β6*gender identity +β7* year in school + β8*pre course difficulty +β9* pre instructor support + □

*Note*. In the above equation, ‘Anxiety’ is a general place-holder for different measures of anxiety across the three models. Model A used mean ESM anxiety score, model B used pre-survey anxiety score, and model C used post-survey anxiety score.

Besides these three models, we also examined how other measures of anxiety predicted final grades, including the standard deviation of students’ ESM responses, average of pre-post measures of anxiety, and the mean level of students’ ESM responses truncated to the first half of the semester. For the sake of brevity, we present the results for these three additional models in the Supplementary (Table S1).

## Results

The results are shown by research question, each aligned with a specific validity standard. Our first research question addresses Standard 1.8 and compares the participant population between ESM (*N* = 176) and non-ESM participants (*N* = 832). For research questions two and three, we focus *solely* on ‘complete observations’ (*N* = 176) and provide validity evidence to address Standards 1.14 and 1.17, respectively (AERA, APA, & NCME, 2014).

### RQ 1a: How do ESM participants compare to non-ESM participants in terms of demographic characteristics?

There were few differences in demographic composition between ESM and non-ESM participants (Table 3). Participants in both groups were majority white, first-year women who had taken a biology class within the past year. Table 3 shows chi-squared statistics and *p*-values for chi-squared tests comparing differences in demographic composition between ESM and non-ESM participants. There were no significant differences between ESM and non-ESM participants in terms of race, year in school, and number of years since students’ last biology class. Two demographic characteristics differed significantly between ESM and non-ESM participants: gender and course type.

**Table 3:** Demographic composition and Chi-squared values comparing participants who participated in ESM and those who did not.

| *Category* | *Sub-group* | *ESM participant count (N= 176)* | *Non-ESM participant count (N = 832)* | $\chi^2$ | *DF* | *p* |
| --- | --- | --- | --- | --- | --- | --- |
| ***Race*** | White | 147 | 671 | 0.608 | 1 | 0.436 |
|  | Non-white | 29 | 161 |  |  |  |
| ***Gender*** | Man | 26 | 196 | 7.027 | 2 | **0.030** |
|  | Woman | 145 | 621 |  |  |  |
|  | Non-binary | 5 | 15 |  |  |  |
| ***Year in*** | 1 | 109 | 531 | 2.580 | 4 | 0.630 |
| *school* | 2 | 48 | 198 |  |  |  |
|  | 3 | 14 | 78 |  |  |  |
|  | 4 | 5 | 19 |  |  |  |
|  | 5 | 0 | 6 |  |  |  |
| *Time since last Biology Class* | 1 | 87 | 434 | 2.997 | 4 | 0.558 |
|  | 2 | 24 | 121 |  |  |  |
|  | 3 | 21 | 114 |  |  |  |
|  | 4 | 31 | 121 |  |  |  |
|  | 5 | 13 | 42 |  |  |  |
| *Course* | Non-majors | 84 | 524* | 73.87 | 2 | <0.001 |
|  | Major<br>s | 92 | 217* |  |  |  |

\* Note: The sample size for non-ESM course calculations is 741, due to missing data for an instructor.

In terms of the gender distribution across ESM and non-ESM participants, standardized residuals of the chi-squared test (χ^2^ = 7.0267, *df* = 2, *p-*value = 0.0298) indicate that there was a significant underrepresentation of men in the ESM group compared to the non-ESM group (Z= – 4.61, *p*-value <0.001).

In terms of course type, chi-squared tests indicated that ESM and non-ESM groups differed significantly (χ^2^ = 7.32.618, *df* = 1, *p-*value < 0.001), with students in the non-majors course significantly overrepresented in our ESM population (Z = 8.14, *p*-value < 0.001) and students in the majors course significantly underrepresented (Z = –5.80, *p*-value <0.001).

### RQ 1b: How do ESM participants compare to non-ESM participants in terms of their pre-course perceptions?

To determine whether either method preferentially recruited participants according to their perceptions (e.g., whether higher anxiety students were more likely to opt-in to ESM, etc.), we examined differences between the two participant groups in terms of students’ perceptions of pre-course anxiety, course difficulty, and instructor support. We found no significant differences between the groups’ mean perceptions of anxiety (W = 74,938, *p* = 0.628) or course difficulty (*W* = 73,737, *p* = 0.882). However, ESM participants had a higher perception of instructor support (*M* = 8.73, *SD* = 1.57) compared to non-ESM participants (*M*= 8.31, *SD* = 1.68), and this difference was statistically significant (*W* = 62042, *p* = 0.001).

### RQ 2: How do data gathered via ESM and pre-post surveys compare in their ability to describe students’ mean level of anxiety and its variability throughout the semester?

The mean level of student anxiety measured via ESM was moderate (*M* = 3.17, *SD* = 1.72). The mean level of student anxiety measured via pre-post surveys was slightly higher (*M* = 3.47, *SD* =1.53). Paired t-tests indicated that mean-levels of anxiety measured via ESM and pre-post surveys differed significantly (t = –4.32, *p* = < 0.001, Figure 4). However, the effect size for this difference was relatively small (Cohen’s *d* = –0.18).

Next, we examined the intra-individual variability in ESM responses to gain a sense for how much respondents’ experiences of anxiety changed throughout the semester. On average, student ESM responses varied by 0.865 Likert-scale-points each time they responded (*M*= 0.865, *SD* = 0.472). Notably, there was substantial variation in this measure, indicating how variable students’ anxiety was over time (Figure 5). We sought to contrast the variation in ESM responses with anxiety measured through pre-post surveys. Because pre-post surveys consist of two measures, we calculated change scores (i.e. post-survey anxiety score minus pre-survey anxiety score), rather than standard deviations, for each individual’s pre-post survey responses. The average change between post– and pre-survey was small (*M* = –0.028, *SD* = 1.59), indicating that, on average, students reported lower anxiety in the post-survey, though there was a large amount of variability between individual students’ change scores.

### RQ 3: How do data gathered via ESM and pre-post surveys compare in terms of their ability to predict end of course outcomes?

Lastly, we sought to compare the predictive ability of data gathered via ESM and pre-post surveys on students’ final course grade. To do this, we ran three models which included the same set of predictors, changing only the measure of anxiety (i.e., mean-level ESM anxiety, pre-survey measure of anxiety, and post-survey measure of anxiety) to test how each related to students’ end of course grade. The output from these models is presented in Table 4, with the critical predictors being the bottom three rows: *Pre-survey Anxiety, Post-survey Anxiety,* and *Mean ESM*, with all other variables being the same except for these different measures of anxiety. The first column contains the unstandardized coefficient and standard error, and the second column contains the standardized coefficient and standard error; because standardizing does not change the *p-*values, these are the same for both coefficients and their standard errors. We use B for the unstandardized coefficients, and β for the standardized coefficients.

**Table 4:**
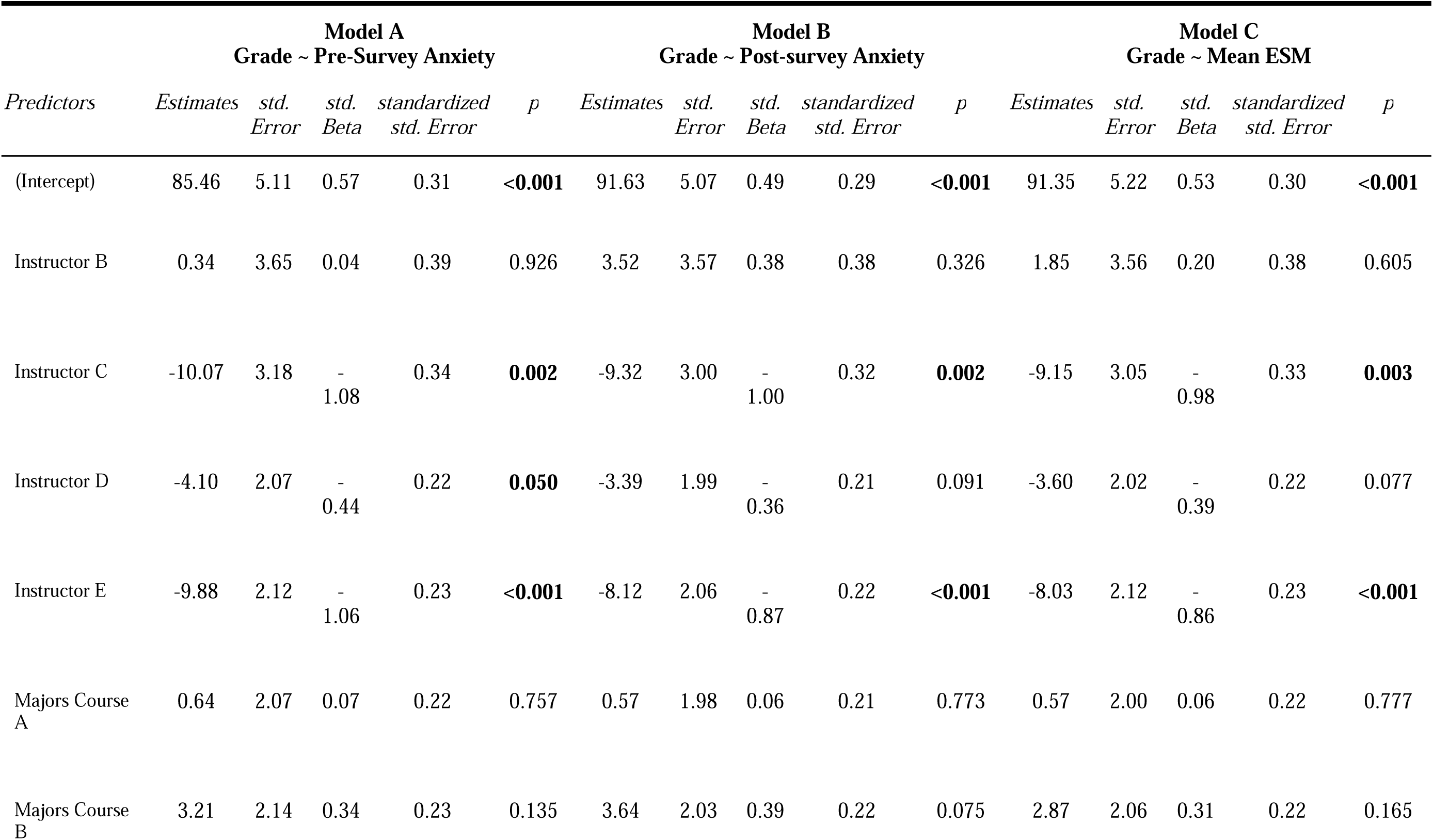
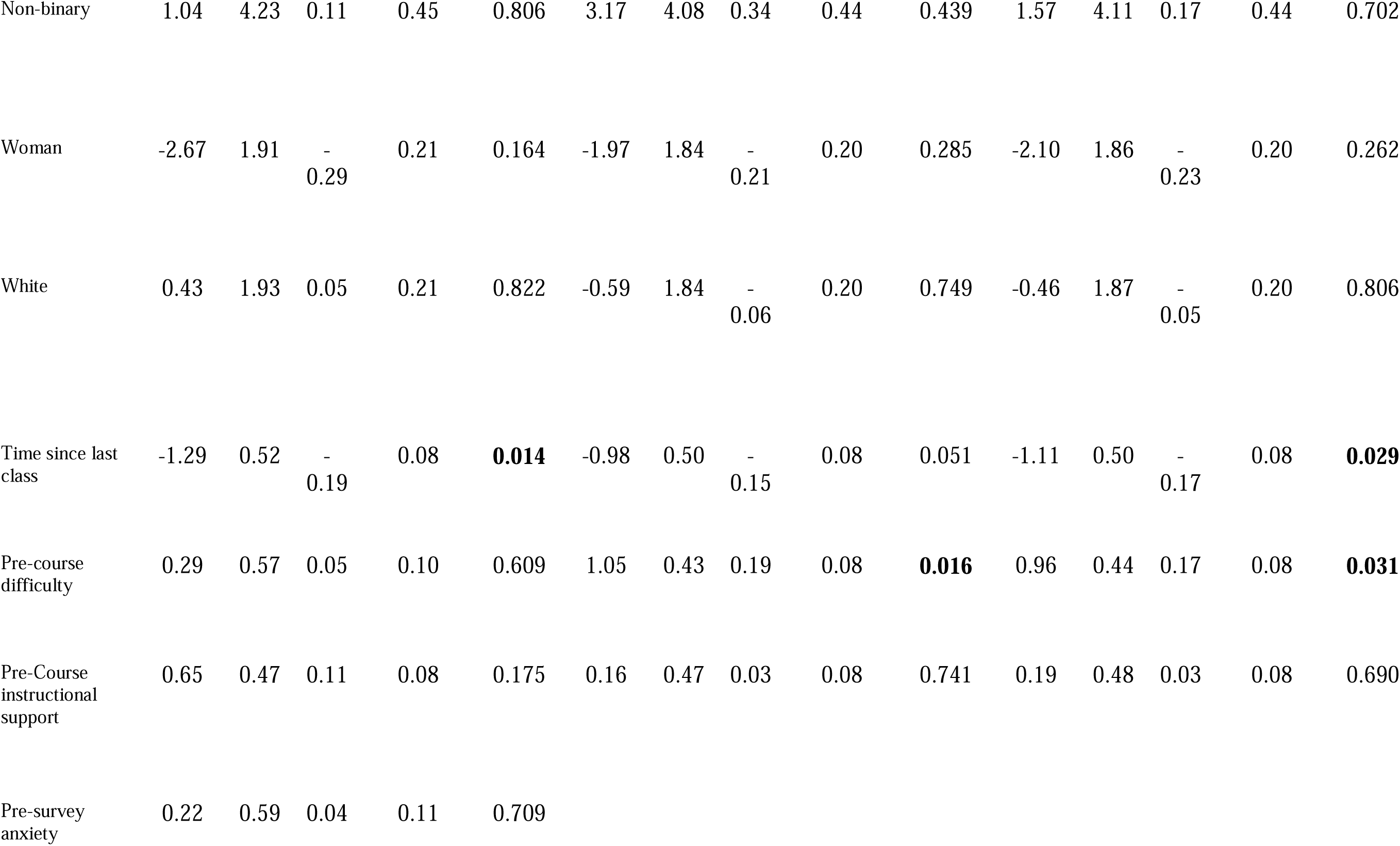
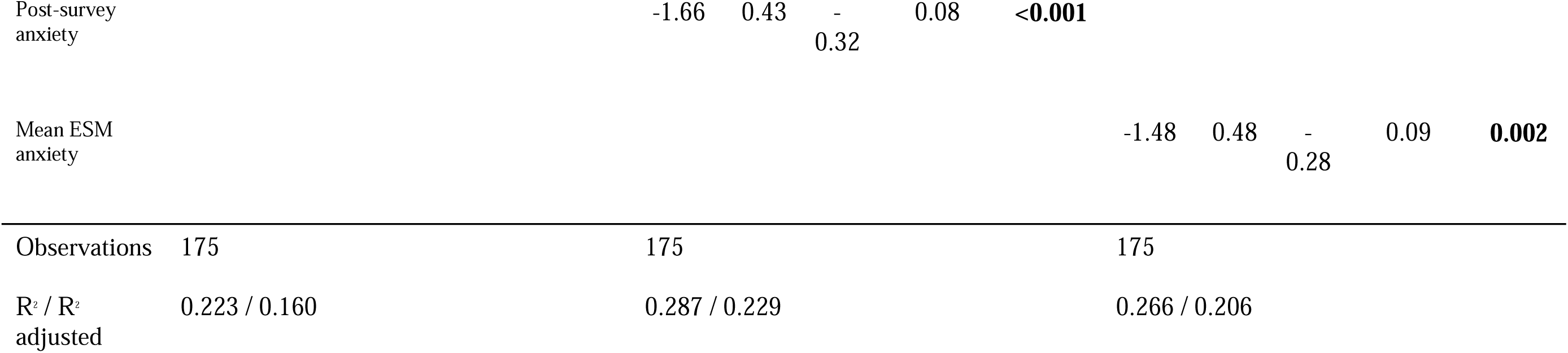
Linear regression output from three models predicting final course grade.

We found no appreciable relationship between pre-survey anxiety and final course grade (B = 0.22 (SE = 0.59), β = 0.04 (SE = 0.11), *p* = .0.709; Table 4). While the coefficient is positive, the standard error is far greater in magnitude, and the *p*-value indicates that this is a highly uncertain estimate of the relation between pre-survey anxiety and grade.

We found a moderate, negative relation between post-survey anxiety and students’ final course grade (B = –1.66 (SE = 0.43), β = –.32 (SE = 0.08), *p* < .001; Table 4). This means that for every one unit increase in students’ post-survey anxiety as measured via an average of their pre– and post-survey anxiety, there is, based upon the model, a –1.66 percentage point decrease in students’ final grade. When both the predictor (i.e., students’ anxiety measured via the average of their pre– and post-survey anxiety) and dependent variable (i.e., students’ final course grade) are both standardized (so that for each *M* = 0, *SD =* 1), there is a –0.32 standard deviation decrease in students’ final grade for each one-standard deviation increase in their mean pre-post anxiety.

We also found a negative relationship between mean ESM and final course grade. For Mean ESM: B = –1.48 (SE = 0.48), β = –.28 (SE = 0.09), *p* = .002. This tells us that for every one-unit increase in students’ mean anxiety as reported through their ESM, there is a –1.48 percentage point decrease in their final grades. On a standardized basis, the coefficient was –.28, there is a –0.28 standard deviation decrease in students’ final grade for each one-standard deviation increase in their mean pre-post anxiety.

## Discussion

Student anxiety is an important aspect of the student experience that can impact students’ achievement, persistence, and well-being. Yet, standard methods used to measure anxiety may be unable to fully capture the nuance that exists within these experiences. As such, this study sought to provide validity evidence for a novel method for measuring anxiety in large introductory biology classrooms: experience sampling methods (ESM). To do this, we compared ESM sampling with traditional pre-post sampling in these classes. We found that ESM and non-ESM participants were relatively similar in terms of demographic and perceptual variables (i.e., pre-course anxiety and difficulty), but differences in the gender distribution of the sample could have implications for anxiety measures. In terms of ESM’s ability to measure student anxiety compared to standard pre-post surveys, we found that, on average, anxiety measured via ESM was lower and the level of variation among individual students’ responses was considerably higher. Finally, we found that pre-survey anxiety was unrelated to students’ final grades, but that average ESM anxiety and the post-survey measures of anxiety were both highly predictive of final grade. Overall, these results provide evidence of validity for the use of ESM to measure anxiety in large, introductory courses, and highlight its unique strengths in capturing variation in student anxiety over time. These results have implications for research measurement and open new avenues for interventions that could ameliorate students’ anxiety and increase their academic success.

We first sought to understand the composition of our ESM sample and how it may have differed from non-ESM participants. The goal of this research question was to provide validity evidence as to whether the samples generated from ESM and traditional pre-post designs are similar and address standard 1.8: “*Describe the composition of any sample in as much detail as possible*” (AERA, APA, & NCME, 2014). Reporting the demographic and perceptual characteristics of a sample is especially important when measuring characteristics, like anxiety, that have strong relationships with individual factors (AERA, APA, & NCME, 2014). Previous work has highlighted gender differences in anxiety, with women in introductory Biology reporting higher average anxiety compared to men (Ballen et al., 2017) as well as an inverse relationship between student perceptions of instructor support and students’ level of anxiety (Schussler et al., 2021; Weatherton et al., *in review*). Therefore, the fact that ESM and non-ESM participants differed significantly in gender, course type, and their perceptions of instructor support may mean that students in our ESM sample had different experiences with anxiety than non-ESM students. As such, our results may not be broadly generalizable to all introductory biology students (see *limitations*). ESM participants’ higher perception of instructor support may underlie more general positive perceptions about their instructor, and thus may have predisposed these students to enroll in additional data collection (i.e., ESM). Previous work has found that women are more likely to enroll in data collection efforts generally, and this, too, may have contributed to the differences among our participant groups (Saleh & Bista, 2017; Marconi et al., 2019). This finding highlights the importance of describing participant populations when using ESM, especially considering that ESM studies have a higher ‘burden’ on participants compared to pre-post methods (Eisele et al., 2022). As such, researchers should be cautious in generalizing any ESM results to the broader population unless sample equivalence can be documented. We suggest that future studies address this validity standard, especially in cases where researchers are interested in factors that are affected by individual differences (e.g., engagement, science identity, etc.), or when analysis methods do not account for individual factors (i.e., when presenting mean-level results as opposed to modeling where demographic factors are accounted for).

To address Standard 1.14, we sought to compare differences in the two methods’ ability to describe students’ mean-level of anxiety and variation in anxiety over time. We found that mean-level anxiety was significantly lower when measured via ESM, and that ESM was able to capture far more variation in students’ anxiety across time. The discrepancy between these measurement tools aligns with previous findings examining student engagement (Xie et al., 2019), and suggest that ESM’s more frequent and real-time sampling may provide more accurate results. These results may be attributed to ESM’s ability to reduce recall and social desirability biases known to affect pre-post survey data (Beal, 2015; Ketonen et al., 2018; Eisele et al., 2022). Moreover, the average anxiety levels obtained via ESM might more accurately reflect the true mean of anxiety, as ESM, by capturing a higher frequency of data points, effectively averages out anxiety’s inherent variability. This contrasts with pre-post surveys, where fewer measurements can lead to a less precise representation of the ‘true mean’ of anxiety levels. The finding that students’ anxiety is highly variable across time is supported by our theoretical framework, Control-Value theory, which posits that student emotions are generated based on students’ perceptions of their control over an achievement-related task and their value for that task (Pekrun, 2006). Typical Biology courses include many different achievement-related tasks like homework assignments, exams, and discussion posts, and it’s likely that each student has different perceptions of control and value for each of these tasks across time. As such, Control-Value theory suggests that student emotions (here, anxiety) would be highly variable over time. The ability to capture this variation in students’ anxiety emerged as a distinct strength of ESM over pre-post surveys and underscores the value of utilizing ESM in the assessment of dynamic constructs, such as student emotion. For example, ESM data in this study revealed specific moments when anxiety levels spiked for some students—insights unattainable with traditional survey methods. Figure 5 shows timepoints where anxiety increased across all students on average, and individual-level data reveal even more details about variation in anxiety across time (Figure 3). This strength of ESM may allow future research to reveal novel insights about student anxiety in introductory Biology; for example, exploring how student perceptions of control or value vary across time, or how intra-individual variation in anxiety impacts student outcomes.

Both post-survey anxiety scores and mean-level ESM anxiety scores significantly predicted students’ final grades, providing validity evidence for the use of ESM-derived anxiety scores for this purpose (Standard 1.18, AERA, APA & NCME, 2014). Previous studies established a link between anxiety and final grades (e.g., Awadalla et al., 2020; England et al., 2017), but our findings extend this knowledge by demonstrating that ESM-derived anxiety scores can predict final grades with accuracy comparable to traditional methods. Although post-survey anxiety emerged as a slightly stronger predictor of final grades than ESM anxiety— evident from the standardized beta estimates in Table 4—this outcome is understandable given that post-survey was administered near the end of the term. However, we found that ESM data were able to predict final grades much earlier in the semester with similar accuracy; specifically, using students’ average ESM anxiety scores from weeks 1 to 8 of the semester proved nearly as effective in predicting final grades as the average ESM anxiety measured across the entire semester, with standardized beta coefficients of –0.25 for mid-semester ESM versus –0.28 for full-term ESM (Table S1). These findings suggest that it may be feasible to use mid-semester ESM anxiety levels to anticipate final grades. Such predictive capability could be beneficial for implementing interventions aimed at reducing student anxiety and improving final course grade. Future research should further explore this possibility. Overall, these results reiterate the negative relationship between anxiety and academic outcomes and highlight the ongoing need to better address student emotions in introductory courses in order to improve academic success.

We believe that our results provide adequate validity evidence for the use of ESM to measure student anxiety in introductory Biology. As such, we now discuss specific applications of ESM and identify situations where it offers advantages over traditional pre-post survey methods. While ESM provides a more nuanced understanding of anxiety, it also demands greater resources and effort than pre-post surveys. Therefore, a cost-benefit analysis becomes essential for researchers considering ESM. Factors to consider include the size and nature of the class, the resources available, and the specific research questions being asked. For instance, pre-post surveys might be more practical in large classes with limited resources, whereas ESM’s detailed insights justify its use in smaller classes or when in-depth analysis of anxiety is crucial. As suggested by Pekrun (2006), intensive longitudinal methods like ESM excel at capturing intra-individual variation, making them particularly effective for research aimed at understanding intra-individual processes, like emotions. Conversely, our data suggest that when the objective is to capture mean-level outcomes, the discrepancy in results between ESM and pre-post surveys is minimal. Future work should investigate potential synergistic use of these methods, such as using ESM for detailed process analysis while employing pre-post surveys for broader outcome evaluations.

We believe that ESM can unveil novel insights into student anxiety experiences, surpassing the capabilities of traditional pre-post surveys. Unlike pre-post surveys, ESM facilitates real-time tracking of anxiety fluctuations and responses to particular events or interventions. This can provide valuable insights into the triggers and patterns of student anxiety, which can inform the development of targeted strategies to reduce anxiety and improve student outcomes. For example, if ESM data reveals that student anxiety spikes around certain assignments or activities, instructors may modify these aspects of the course to make them less anxiety-inducing. Similarly, if ESM data show that certain students experience consistently high levels of anxiety, additional support or resources can be provided to these students. ESM may also be used to further refine our theoretical understanding of student emotions; for example, future work may use ESM to evaluate how students’ control and value beliefs vary over time, and how that variation impacts their affective outcomes. Thus, while ESM requires more effort to implement, it has the potential to greatly enhance our understanding of student anxiety, improve the efficacy of educational interventions, and increase students’ success in critical gateway courses like introductory Biology.

## Limitations

Our study is subject to several limitations that warrant mention. First, the generalizability of our findings may be constrained due to the composition of our ESM sample, which was different from both non-ESM participants and students who did not participate in the study at all (see Table S2). The incentive structure for participation, which included bonus points for survey completion and payment for ESM participation, may have additionally influenced the makeup of our participant pool. Furthermore, our data collection spanned only a single semester, limiting our ability to detect differences in student anxiety outside of this timeframe. The number of different courses that were included in our sample prevented us from systematically examining or modeling high anxiety moments throughout the semester, as each course had its own exam schedules and assignment due dates. Finally, it is important to acknowledge the inherent subjectivity of anxiety as a personal experience, which may lead to variability in how students perceive and report their levels of anxiety, potentially influencing the study’s findings. Overall, while our study provides valuable insights into the relationship between anxiety levels and academic performance, these limitations highlight the need for cautious interpretation of the results and underscore the importance of further research to confirm and expand upon our findings.

## Conclusion

Given the scarcity of ESM applications within large, introductory Biology classes, this study sought to establish robust validity evidence for the use of ESM that underscores its efficacy and accuracy in capturing the dynamic landscape of anxiety. Furthermore, this study sought to compare the validity evidence of ESM with that of traditional survey methods, in order to understand the unique strengths and costs of each method. Overall, this study found that ESM and pre-post surveys provided similar results, though there were differences in the details revealed between the methods. These insights not only contribute to our theoretical understanding of student emotions but also provide guidance to educators and researchers aiming to mitigate anxiety and enhance student outcomes. The findings from this research encourage a thoughtful consideration of method selection based not on convention but on the specific aims and constraints of ESM and pre-post surveys in each study.

---

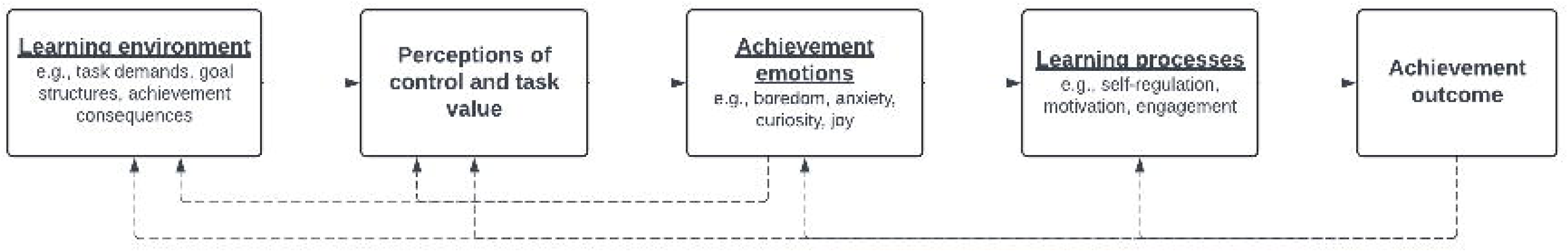

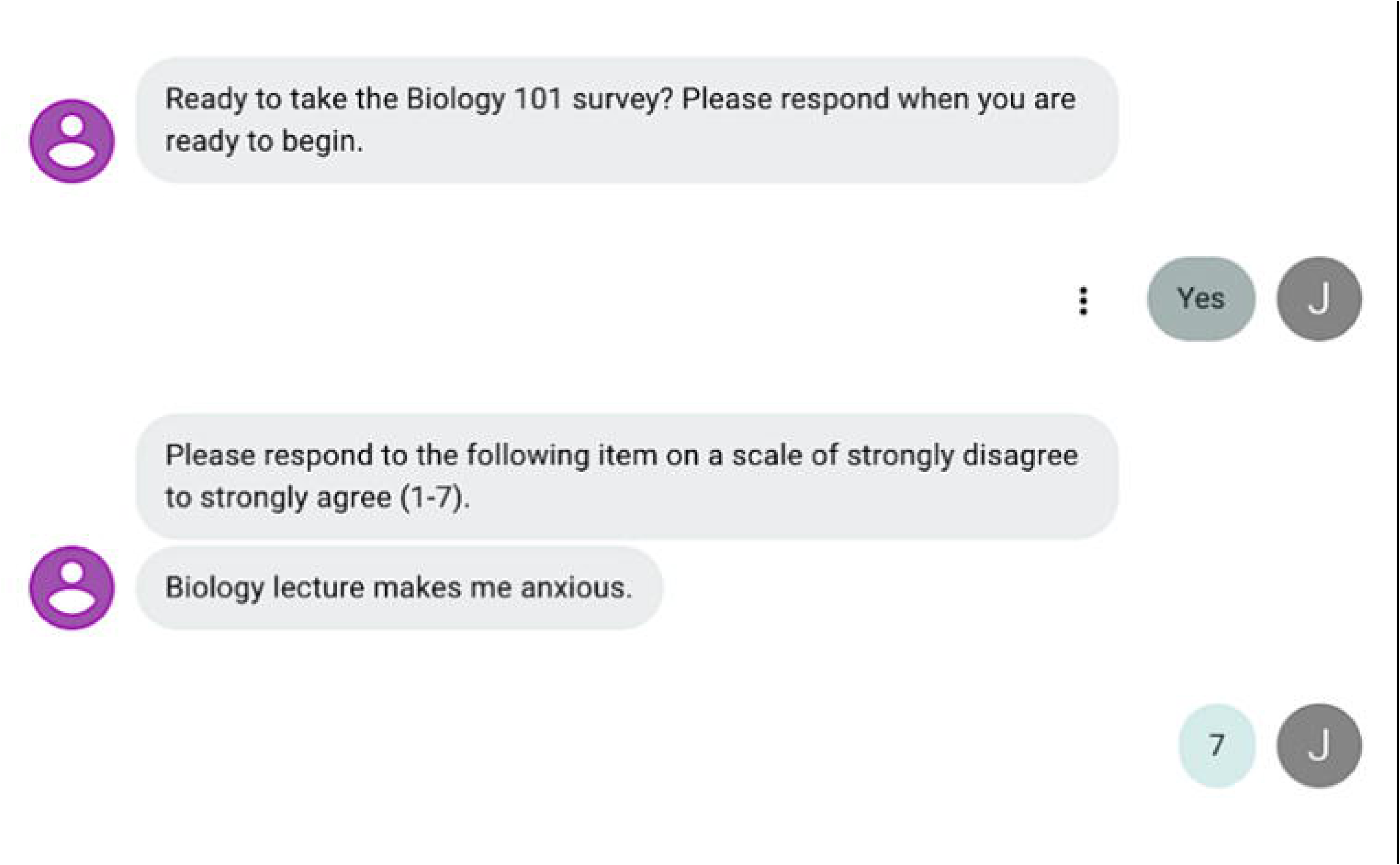

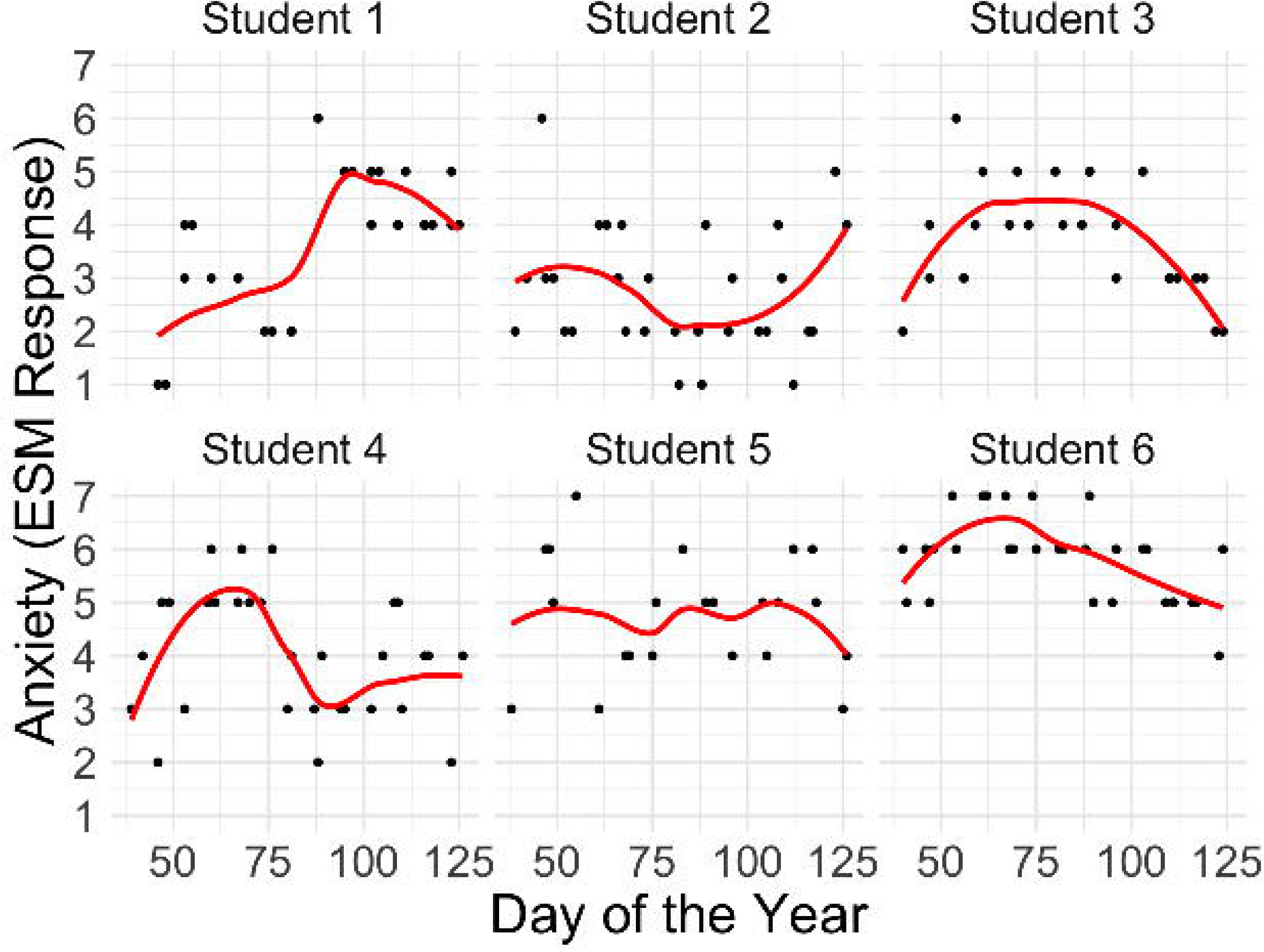

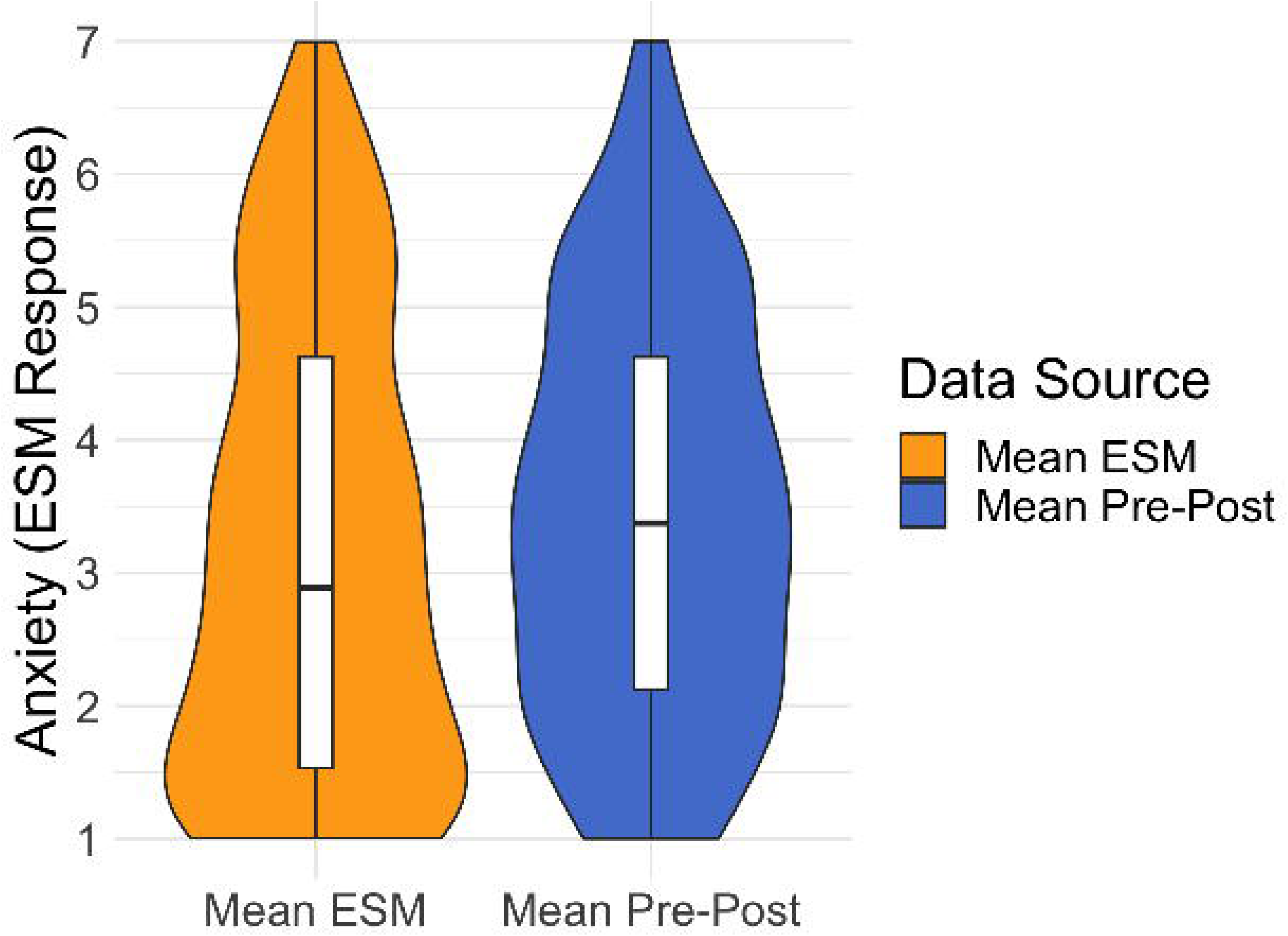

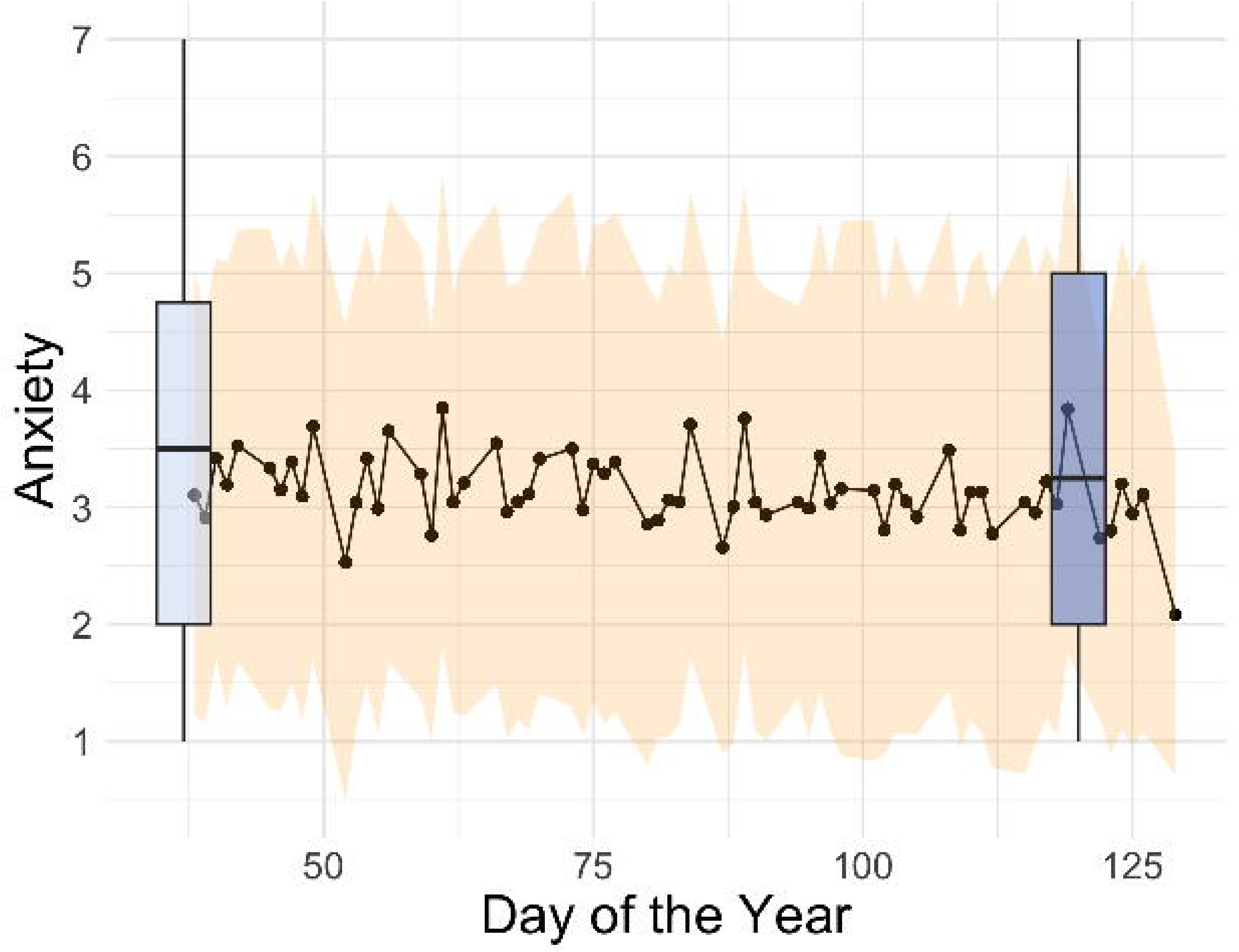

