## Supplementary Appendix B for "Beyond Pre-Post Surveys: Exploring Validity Evidence For The Use Of Experience Sampling Methods To Measure Student Anxiety In Introductory Biology"

Appendix B: Additional Results

**Table S1: Demographic Comparison between ESM participants, non-ESM participants (i.e., those who only completed pre and post survey), and overall course enrollment from Registrar data.**

|  | ESM % (N=176) | Non-ESM % (N=832) | Registrar % (N=1884) |
| --- | --- | --- | --- |
| RACE* |  |  |  |
| White | 83.52 | 80.65 | 75.32 |
| Black or African American | 5.32 | 6.07 | 6.58 |
| Two or more races | 4.22 | 4.37 | 5.52 |
| Hispanic or Latino or Spanish Origin of any race | 3.22 | 4.06 | 7.43 |
| Asian | 3.19 | 3.40 | 3.61 |
| Open Response | 0.53 | 1.21 | -- |
| American Indian or Alaskan Native | 0 | 0.12 | -- |
| Native Hawaiian or Other Pacific Islander | 0 | 0.12 | 0.05 |
| GENDER** |  |  |  |
| Woman | 82.39 | 74.64 | 69.75 |
| Man | 14.77 | 23.56 | 30.25 |
| Non-Binary | 2.31 | 1.19 | -- |
| Prefer to self-describe | 0.53 | 0.61 | -- |
| YEAR IN PROGRAM | |  |  |
| 1 | 61.93 | 63.82 | 53.56 |
| 2 | 27.27 | 23.80 | 30.63 |
| 3 | 7.95 | 9.38 | 11.89 |
| 4 | 2.84 | 2.28 | 3.72 |
| >4 | 0 | 0.72 | 0.05 |
| COURSE |  |  |  |
| Majors’ | 52.27 | 29.28*** | 59.34 |
| Non-Majors’ | 47.73 | 70.72*** | 40.66 |
| *Note that institutional data collects racial categories according to IPEDS categorizations, and therefore does not match up exactly with the way researchers asked about race | | | |
| **Note that institutional data only offers two choices for gender identity, Male or Female. We aligned these choices with ‘Man’ and ‘Woman’, respectively, in this table.    ***Note that the sample size for this calculation is 741, due to data missingness from one course section | | | |

**Table S2: Linear regression output from three additional models predicting final course grade.**

|  | **Model D**  **Grade ~ Mean ESM Anxiety (Week 1:6)** | | | | | **Model E**  **Grade ~ SD (ESM Anxiety)** | | | | | **Model F**  **Grade ~ Average Pre-Post Anxiety** | | | | |
| --- | --- | --- | --- | --- | --- | --- | --- | --- | --- | --- | --- | --- | --- | --- | --- |
| *Predictors* | *Estimates* | *std. Error* | *std. Beta* | *standardized std. Error* | *p* | *Estimates* | *std. Error* | *std. Beta* | *standardized std. Error* | *p* | *Estimates* | *std. Error* | *std. Beta* | *standardized std. Error* | *p* |
| (Intercept) | 91.06 | 5.29 | 0.56 | 0.30 | **<0.001** | 85.41 | 5.36 | 0.57 | 0.31 | **<0.001** | 89.27 | 5.18 | 0.56 | 0.30 | **<0.001** |
| Instructor B | 1.25 | 3.57 | 0.13 | 0.38 | 0.727 | 0.44 | 3.64 | 0.05 | 0.39 | 0.903 | 2.09 | 3.64 | 0.22 | 0.39 | 0.567 |
| Instructor C | -9.68 | 3.06 | -1.04 | 0.33 | **0.002** | -9.81 | 3.15 | -1.05 | 0.34 | **0.002** | -8.99 | 3.10 | -0.97 | 0.33 | **0.004** |
| Instructor D | -4.07 | 2.03 | -0.44 | 0.22 | **0.047** | -4.07 | 2.08 | -0.44 | 0.22 | 0.051 | -3.73 | 2.05 | -0.40 | 0.22 | 0.070 |
| Instructor E | -8.44 | 2.11 | -0.91 | 0.23 | **<0.001** | -9.81 | 2.12 | -1.05 | 0.23 | **<0.001** | -8.70 | 2.12 | -0.94 | 0.23 | **<0.001** |
| Majors Course A | 0.36 | 2.02 | 0.04 | 0.22 | 0.859 | 0.58 | 2.06 | 0.06 | 0.22 | 0.780 | 0.35 | 2.03 | 0.04 | 0.22 | 0.862 |
| Majors Course B | 2.77 | 2.07 | 0.30 | 0.22 | 0.182 | 3.06 | 2.12 | 0.33 | 0.23 | 0.151 | 2.97 | 2.08 | 0.32 | 0.22 | 0.155 |
| Non-binary | 1.85 | 4.14 | 0.20 | 0.45 | 0.656 | 1.05 | 4.24 | 0.11 | 0.46 | 0.805 | 2.27 | 4.19 | 0.24 | 0.45 | 0.589 |
| Woman | -2.01 | 1.88 | -0.22 | 0.20 | 0.287 | -2.67 | 1.92 | -0.29 | 0.21 | 0.166 | -2.22 | 1.89 | -0.24 | 0.20 | 0.241 |
| White | -0.39 | 1.88 | -0.04 | 0.20 | 0.836 | 0.37 | 1.92 | 0.04 | 0.21 | 0.849 | -0.45 | 1.90 | -0.05 | 0.20 | 0.813 |
| Time since last class | -1.15 | 0.51 | -0.17 | 0.08 | **0.024** | -1.25 | 0.52 | -0.19 | 0.08 | **0.018** | -1.06 | 0.52 | -0.16 | 0.08 | **0.040** |
| Pre-course difficulty | 0.93 | 0.45 | 0.17 | 0.08 | **0.039** | 0.41 | 0.43 | 0.07 | 0.08 | 0.340 | 1.17 | 0.51 | 0.21 | 0.09 | **0.025** |
| Pre-Course instructional support | 0.26 | 0.48 | 0.04 | 0.08 | 0.596 | 0.64 | 0.48 | 0.11 | 0.08 | 0.180 | 0.37 | 0.48 | 0.06 | 0.08 | 0.435 |
| **Mean ESM Anxiety halfway** | -1.38 | 0.50 | -0.25 | 0.09 | **0.007** |  |  |  |  |  |  |  |  |  |  |
| **Standard Deviation ESM Anxiety** |  |  |  |  |  | 0.31 | 1.57 | 0.02 | 0.08 | 0.844 |  |  |  |  |  |
| **Mean Pre-Post Anxiety** |  |  |  |  |  |  |  |  |  |  | -1.43 | 0.61 | -0.24 | 0.10 | **0.020** |
| Observations | 175 | | | | | 175 | | | | | 175 | | | | |
| R^2^ / R^2^ adjusted | 0.257 / 0.197 | | | | | 0.222 / 0.159 | | | | | 0.248 / 0.187 | | | | |
