## Supplementary Appendix A for "Beyond Pre-Post Surveys: Exploring Validity Evidence For The Use Of Experience Sampling Methods To Measure Student Anxiety In Introductory Biology"

Spring 2022 Pre-Survey

Start of Block: Default Question Block

Q1
This survey is part of a research study investigating student anxiety* in Biology classes. The questions will be asking about your Biology LECTURE class this semester, specifically the levels of anxiety you may feel during class lecture.

*Anxiety and stress are different, but if you are feeling stress, you are probably also feeling anxiety.

You have logged in to confirm your identity to the researchers. After reading this consent statement, you will be asked whether you consent for them to use your responses as part of their research. If you select “no,” you will still receive any points that your instructor is offering, but your responses will not be a part of the data set for this project. You will also be asked whether you consent to the researchers requesting your final course grade from your instructor after the semester is over. If you say yes, this grade will be matched with your survey responses, stored on a password-protected computer, and accessible only to the researchers.

This project will collect responses at the beginning and end of the semester. We are asking for your names on the survey so we can link your responses to both surveys and then your final course grade. The researchers will also be retaining your responses with your name so that if you respond to this same survey in a future biology class, your responses can be matched. Your name would never be reported along with your responses in any published results.

The survey is voluntary and asks you to answer a variety of questions, as well as provide demographic information. It may take 5-10 minutes to complete.

Your responses will not be "graded" on the basis of your responses. Any points given as an incentive will be based on completion of the survey and not particular responses. If your instructor has decided to award you points for completion of this survey, you will receive those points at the end of the semester, before the final exam. Should you choose to not complete this survey, an alternate assignment will be made available to you so that you can earn the equivalent number of points; this assignment will require the same length of time and effort as required by this survey. Please let your instructor know if you would like this option.

Your instructor will not see the results of this survey until after grades have been turned in; if your instructor chooses to view the responses they will be made anonymous before your instructor would view them. Therefore, the risk associated with your completing this survey is low. The IP address of your computer will not be recorded. However, all internet surveys have the potential for responses to be intercepted; therefore, we cannot guarantee confidentiality.

There is no direct benefit to individuals completing the survey. However, survey responses will be analyzed to determine levels of anxiety currently experienced in Biology lecture classes. Based on respondents’ qualitative comments on this survey, the researchers hope to determine what coping mechanisms might be useful for successfully managing anxiety in a healthy way. The researchers also may take note of student-reported instructor practices that may be causing students anxiety. The elucidation of coping mechanisms for dealing with anxiety as well as instructor practices that could be causing anxiety may be useful for future STEM students, improving their retention rates, which could lead to a larger and more diverse STEM community.

We will not keep your information to use for any future research project. We will also not share your research data with other researchers.

Q2 May we use your responses as part of our research?

- Yes (1)
- No (2)

Q3 May we ask for your final course grade from your course instructor AFTER the semester is over?

- Yes (1)
- No (2)

Q4 Are you age 18 or above?

- Yes (1)
- No (2)

| Page Break |
| --- |

Q5 Please respond to the following items on a scale of strongly disagree to strongly agree.

|  | Strongly disagree (1) | Disagree (2) | Somewhat disagree (3) | Neither agree nor disagree (4) | Somewhat agree (5) | Agree (6) | Strongly agree (7) |
| --- | --- | --- | --- | --- | --- | --- | --- |
| Biology lecture makes me nervous (1) |  |  |  |  |  |  |  |
| Biology lecture is stressful (2) |  |  |  |  |  |  |  |
| Biology lecture makes me anxious (3) |  |  |  |  |  |  |  |
| Biology lecture scares me (4) |  |  |  |  |  |  |  |
| Biology lecture is complex (5) |  |  |  |  |  |  |  |
| Biology lecture is complicated (6) |  |  |  |  |  |  |  |
| Biology lecture is difficult (7) |  |  |  |  |  |  |  |

Q6 Why do you feel this way about your biology lecture this semester?

________________________________________________________________

| Page Break |
| --- |

Q7 Answer:

|  | 1 - not supportive (1) | 2 (2) | 3 (3) | 4 (4) | 5 (5) | 6 (6) | 7 (7) | 8 (8) | 9 (9) | 10 - very supportive (10) |
| --- | --- | --- | --- | --- | --- | --- | --- | --- | --- | --- |
| On a 10-point scale, how supportive do you feel the instructor of your biology class has been so far? (1) |  |  |  |  |  |  |  |  |  |  |

Q8 What specific things has your instructor SAID or DONE that led to the support rating you gave them? (if you remember these specifics)

________________________________________________________________

Q9 Besides specific things your instructor said or did, what else informed your rating of their supportiveness?

________________________________________________________________

Q10 There are no "right" or "wrong" answers. Your opinion is what is wanted on each item. Please think about how well each statement describes what this course is like for you using the response categories always, often, sometimes, seldom, or never. In this class...

|  | Always (1) | Often (2) | Sometimes (3) | Seldom (6) | Never (4) |
| --- | --- | --- | --- | --- | --- |
| If I have an inquiry, the instructor finds time to respond. (1) |  |  |  |  |  |
| The instructor helps me identify problem areas in my study. (20) |  |  |  |  |  |
| The instructor responds promptly to my questions. (21) |  |  |  |  |  |
| The instructor gives me valuable feedback on my assignments. (22) |  |  |  |  |  |
| The instructor adequately addresses my questions. (23) |  |  |  |  |  |
| The instructor encourages my participation. (24) |  |  |  |  |  |
| It is easy to contact the instructor. (25) |  |  |  |  |  |
| The instructor provides me with positive and negative feedback on my work. (26) |  |  |  |  |  |

| Page Break |
| --- |

Q11 Choose answer choice 2 for this question.

|  | 1 (1) | 2 (2) | 3 (3) | 4 (4) | 5 (5) |
| --- | --- | --- | --- | --- | --- |
| This question is asking you to choose the number 2. (1) |  |  |  |  |  |

| Page Break |
| --- |

Q12 What year are you in school?

- 1 (freshman) (1)
- 2 (2)
- 3 (3)
- 4 (4)
- >4 (5)

Q13 What is your gender identity?

- Woman (1)
- Man (2)
- Non-binary (4)
- Prefer to self-describe below (6) __________________________________________________

Q14 What is your racial/ethnic identity?

- Hispanic or Latino or Spanish Origin of any race (1)
- American Indian or Alaskan Native (2)
- Asian (3)
- Native Hawaiian or Other Pacific Islander (4)
- Black or African American (5)
- White (6)
- Two or more races (7)
- Open Response (8) __________________________________________________

| Page Break |
| --- |

Q15 What is your current or intended major?

________________________________________________________________

Q16 What is the name of the professor of your Biology lecture class this semester? [QUESTION HAS BEEN BLINDED FOR INSTRUCTOR PRIVACY]

- (1)
- (2)
- (3)
- (4)
- (5)
- (6)
- (7)

Q17 How long has it been since you last took a biology class?

- 0-1 years (1)
- 1-2 years (2)
- 2-3 years (3)
- 3-4 years (4)
- more than 4 years (5)

| Page Break |
| --- |

Q18 If you have additional thoughts about your Biology class, please write them below.

________________________________________________________________

________________________________________________________________

________________________________________________________________

________________________________________________________________

________________________________________________________________

Q19 Please type the following here: Last name

________________________________________________________________

________________________________________________________________

________________________________________________________________

________________________________________________________________

________________________________________________________________

Q20 Please type the following here: First name

________________________________________________________________

Q21 Please type the following here: Net ID (e-mail)

________________________________________________________________

| Page Break |
| --- |

Q22
We are seeking participants to explore the use of text messages as survey responses.

Would you like information about participating in the text messaged-based study?

You will be compensated (with an Amazon.com gift card valued up to $30) for participating in the text messaged-based portion of the study.

- Yes (this option will complete the survey and take you to information about the text message-based study) (1)
- No (this option will complete the survey) (2)

End of Block: Default Question Block
